# De novo design of protease-activatable cytokine prodrugs

**DOI:** 10.64898/2026.09.19.752834

**Authors:** Hojeong Shin, Julissa G. Tello, Chan Johng Kim, Jung-Ho Chun, Frank Peprah, Brendan D. Parent, Tavus Atajanova, Robiah Arefin Ibn Mahmud, Li Qiang, Andrew J. Aguirre, Stephanie K. Dougan, Michael Dougan, David Baker

## Abstract

Cytokines can elicit potent antitumor immunity, but their clinical use is often limited by systemic immune activation and dose-limiting toxicity. Protease-responsive masking offers a way to improve selectivity by linking cytokine activity to disease-associated protease cleavage, but current approaches have faced several challenges. Here we describe a de novo protein design strategy for generating protease-activatable cytokine prodrugs that overcomes these challenges. We use this strategy to design and characterize cytokine prodrugs and well-behaved AND-gated split systems that require both target-dependent colocalization and proteolytic unmasking to generate active cytokines. In challenging syngeneic tumor models, our IL-21 prodrugs reduced treatment-associated toxicity while maintaining antitumor efficacy.

## Introduction

Cytokines function as both regulators and effectors of antitumor immunity, shaping immune-cell proliferation, differentiation, and activity, and are therefore attractive agents for cancer immunotherapy^1–3^. Their therapeutic use, however, is often limited by these same properties: systemic exposure can activate cytokine-responsive cells outside the tumor, producing a narrow therapeutic window and dose-limiting toxicity^1,4,5^. This problem extends beyond native cytokines to engineered agonists and de novo-designed cytokine mimics, which can retain strong antitumor activity but still activate receptors in peripheral tissues when administered systemically^6,7^. Several strategies have been developed to restrict cytokine activity to diseased tissues^4^, including tumor targeting^8^, affinity attenuation^9^, split cytokine architectures^10^, and protease-responsive masking^11–13^. Protease-responsive masking is particularly appealing because it links cytokine activation to enzymatic activities enriched in pathological microenvironments^14,15^. However, successful cytokine masking depends critically on the coupled properties of the masking domain and protease-cleavable linker. The mask must engage the cytokine with an affinity and geometry that suppress receptor binding in the intact prodrug, without preventing activity recovery after cleavage. Linker cleavage must likewise permit efficient local activation while avoiding premature systemic release of active cytokine. These constraints are difficult to satisfy using native receptor-derived ectodomains or naturally occurring inhibitory partners, whose size, affinity, and binding geometry are largely fixed.

We reasoned that de novo protein design could overcome these constraints by creating compact masking domains tailored to the receptor-binding surfaces of cytokine mimics. In this strategy, the mask is built to sterically occlude receptor engagement in the intact prodrug, and protease cleavage restores activity by separating the mask from the cytokine mimic (Fig. 1a). We refer to these de novo-masked, protease-activatable cytokine mimics as Prokines. Because the mask is designed directly from the cytokine structure rather than borrowed from a native receptor, its placement, size, and affinity can be tuned independently, enabling alternative mask designs to be explored and optimized for each cytokine scaffold. We set out to use this approach to develop IL-21- and IL-2-like prodrugs and AND-gated split cytokine systems in which activity requires both proteolytic unmasking and target-dependent reconstitution.

**Figure 1.**
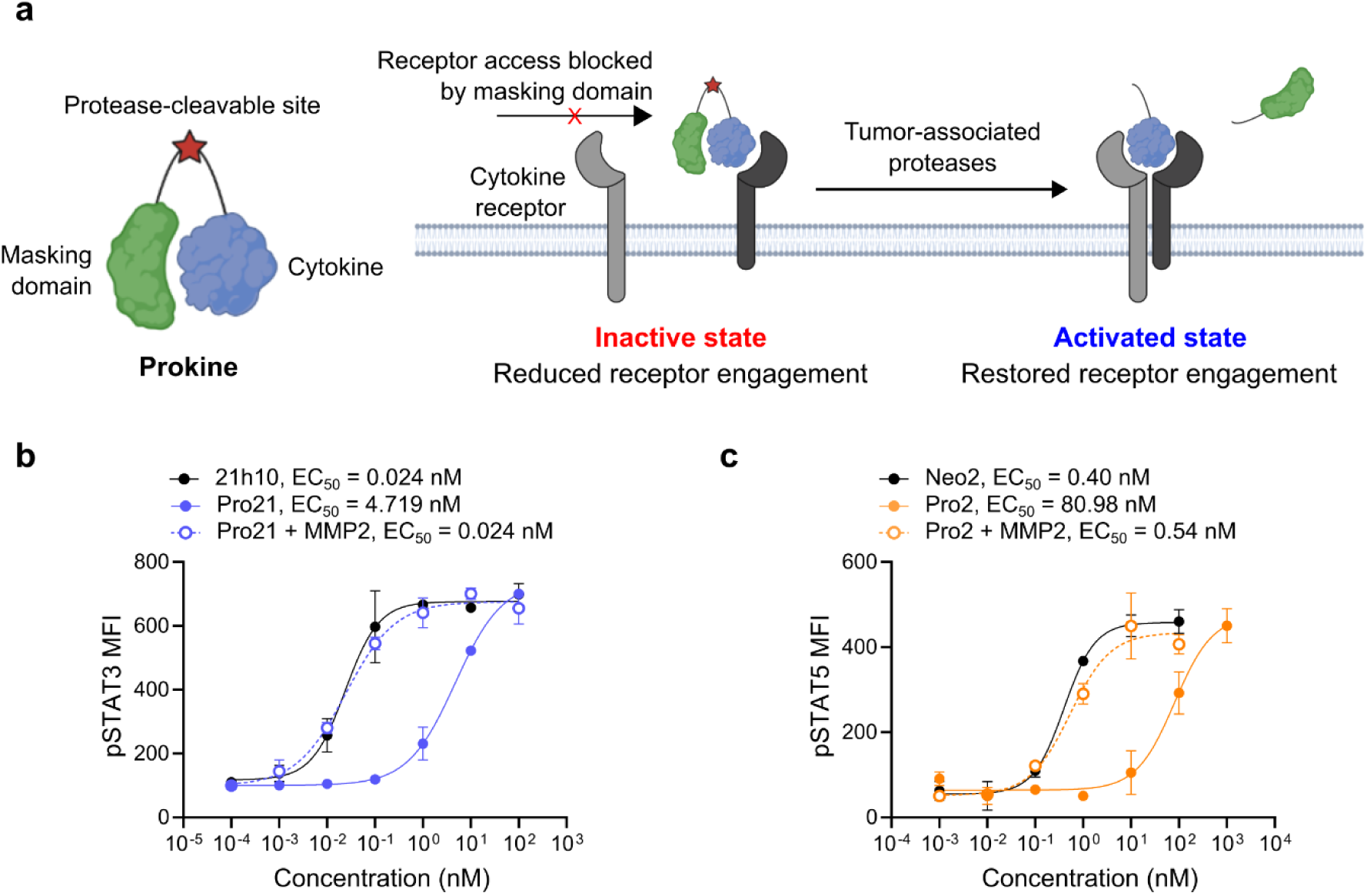
De novo design of protease-activatable IL-21 and IL-2 mimics. **a,** Schematic of the Prokine architecture. A de novo-designed masking domain is connected to a cytokine mimic through a protease-cleavable linker and limits receptor engagement in the intact state. Proteolytic cleavage releases the mask and restores access to the receptor-binding surface. **b,c,** Human PBMC signaling assays measuring pSTAT3 induction by 21h10, Pro21, and MMP2-cleaved Pro21 (**b**) or pSTAT5 induction by Neo2, Pro2, and MMP2-cleaved Pro2 (**c**). Pro21 and Pro2 showed reduced signaling in the intact masked state, whereas MMP2 treatment restored signaling to near-parental potency. EC50 values are indicated in each panel. Data are mean ± s.d. of three technical replicates.

## Results

### Structure-guided design of protease-activatable cytokine prodrugs

We first sought to determine whether de novo-designed proteins could serve as masks for cytokine mimics. We selected the IL-21 mimic 21h10 and the IL-2 mimic Neo2 as potent and structurally characterized agonists that are cross-reactive with the corresponding human and murine receptor complexes^6,7^. For each cytokine mimic, residues within the receptor-binding interface were specified as design hotspots, with the goal of generating masks that occlude receptor engagement. We used RFdiffusion2^16^ to generate structurally distinct candidate masking backbones at these sites, ProteinMPNN^17^ for sequence design, and AlphaFold2 (AF2) initial guess predictions^18^ to prioritize high-confidence mask–cytokine complexes for experimental testing.

Selected candidates were fused to MMP2- and MMP9-sensitive linkers, because both proteases are frequently elevated in tumor microenvironments and have conserved orthologues in mice and humans^19–21^. The resulting Prokines were screened by biolayer interferometry (BLI) and cell-based signaling assays. The initial IL-21 and IL-2 leads, D10 and A3, respectively, attenuated signaling in human peripheral blood mononuclear cells (PBMCs) while retaining MMP-dependent reactivation. In CD3-positive cells, D10 produced an approximately 17-fold shift in 21h10 potency between the intact and MMP-treated states (Fig. S1a), and A3 produced an approximately 16-fold shift in Neo2 activity (Fig. S1b). These results established that de novo masks could suppress cytokine signaling and that activity could be recovered by proteolysis of the linker, but the residual signaling in the intact state indicated that stronger receptor blockade was needed. We used interface-focused ProteinMPNN redesign to increase masking strength, retaining the overall mask backbone while selectively diversifying residues at the mask–cytokine interface. This yielded optimized IL-21 and IL-2 Prokines, termed Pro21 and Pro2, respectively.

Structural models suggested that the masks occupy the intended receptor-binding regions in both constructs, consistent with the design goal of suppressing signaling by blocking access to the receptor interface. Both Pro21 and Pro2 were expressed as soluble proteins with predominant size-exclusion chromatography peaks, whereas the corresponding native receptor-masked constructs were recovered at substantially lower levels following expression and purification from cultures of the same expression scale (Fig. S2a,b). Circular dichroism showed that both Pro21 and Pro2 were highly structured across a broad temperature range, with melting temperatures of >95 °C and ∼88 °C, respectively (Fig. S3a–d). Surface plasmon resonance confirmed that masking strongly reduced receptor binding. In the IL-21 system, 21h10 bound IL-21R with a K_D_ of 21.8 pM, whereas Pro21 binding was reduced to 15.1 μM (Fig. S4a). In the IL-2 system, Neo2 bound IL-2Rβ with a K_D_ of 4.41 nM, whereas Pro2 binding was reduced to 748 nM (Fig. S4b). Thus, structure-guided design and interface-focused optimization generated compact de novo masks that strongly attenuate receptor binding while preserving favorable biophysical properties.

### Proteolytic unmasking restores cytokine signaling and enables targeted activation

We next verified that the optimized Prokines were proteolytically processed by the intended proteases. MMP2 and MMP9 treatment generated cleavage products corresponding to the cytokine mimic and masking domain for Pro21 and Pro2, whereas the corresponding non-cleavable (NC) controls remained predominantly intact (Fig. S5a,b). In human PBMCs, Pro21 and Pro2 showed substantially attenuated signaling in the masked state, with EC50 values shifted by approximately 200-fold relative to their parental cytokine mimics. MMP2 treatment restored signaling to near-parental potency (Fig. 1b,c).

We then asked whether protease-activatable masking could be combined with a second layer of spatial control through target-cell localization. We tested PD-L1-targeted Pro21 in PBMC co-cultures containing either wild-type B16 cells or PD-L1-overexpressing B16 cells. PD-L1-targeted Pro21 preferentially activated PBMCs in the presence of PD-L1-overexpressing tumor cells (Fig. S6). Together, these results showed that Pro21 and Pro2 function as MMP-activatable cytokine prodrugs whose signaling can be restored by proteolytic unmasking. PD-L1 targeting further biased Pro21 activity toward antigen-expressing target cells.

### De novo masking enables protease- and tumor-antigen-dependent AND-gated cytokine activation

We investigated whether de novo masking could address a related but distinct challenge in split cytokine design. Splitting a cytokine mimic provides a route to conditional activation because signaling requires productive assembly of two fragments. However, each fragment can be poorly behaved on its own, complicating drug development, and may reconstitute before reaching the intended site of action, resulting in background activity. In our previous work, the two split Neo2 components were administered at separate injection sites to avoid premature reconstitution^10^. We reasoned that masks designed against the fragment interfaces required for reconstitution could suppress premature assembly and convert split Neo2 into a two-input system requiring both protease cleavage and target-dependent colocalization.

Neo2 was divided into two fragments, Neo2a and Neo2b, whose combined signaling activity depends on reconstitution. Using the design framework described above for masking intact cytokine mimics, we generated masked fragments Pro2a and Pro2b in which de novo masks were positioned to block the surfaces required for fragment assembly. This strategy also resolves a biophysical challenge created by splitting: isolated fragments can expose hydrophobic surfaces that are normally buried in the intact cytokine. Consistent with this, unmasked Neo2a showed little detectable monodisperse peak by size-exclusion chromatography, whereas Pro2a eluted as a predominant monodisperse species. Pro2b also eluted as a well-defined monodisperse peak (Fig. S7a).

Functional assays showed that the masks imposed strong protease control over split cytokine activity. In HEK-Blue IL-2 reporter cells, Neo2a and Neo2b reconstituted signaling with an EC50 of 1.216 nM. In contrast, the Pro2a and Pro2b pair had no measurable EC50 up to 1 µM, the highest concentration tested, corresponding to at least 800-fold attenuation relative to the unmasked split pair (Fig. S7b). MMP2 treatment restored activity to near that of the unmasked split pair.

Because activity of the split cytokine requires assembly of two components, we reasoned that it should depend on effective local concentration on antigen-expressing cells. To test this, both split components were fused to a PD-L1 minibinder and assayed in co-cultures with wild-type or PD-L1-overexpressing B16 cells (Fig. 2a). The unmasked targeted pair activated reporter cells selectively in the presence of PD-L1-overexpressing cells, whereas substantially lower activity was observed with wild-type cells. In contrast, the masked targeted pair showed little activity even in the presence of PD-L1-overexpressing cells prior to MMP2 treatment, indicating that antigen-dependent colocalization alone was not sufficient to activate the masked split cytokine. After MMP2 cleavage, activity was restored selectively in the presence of PD-L1-overexpressing cells (Fig. 2b).

**Figure 2.**
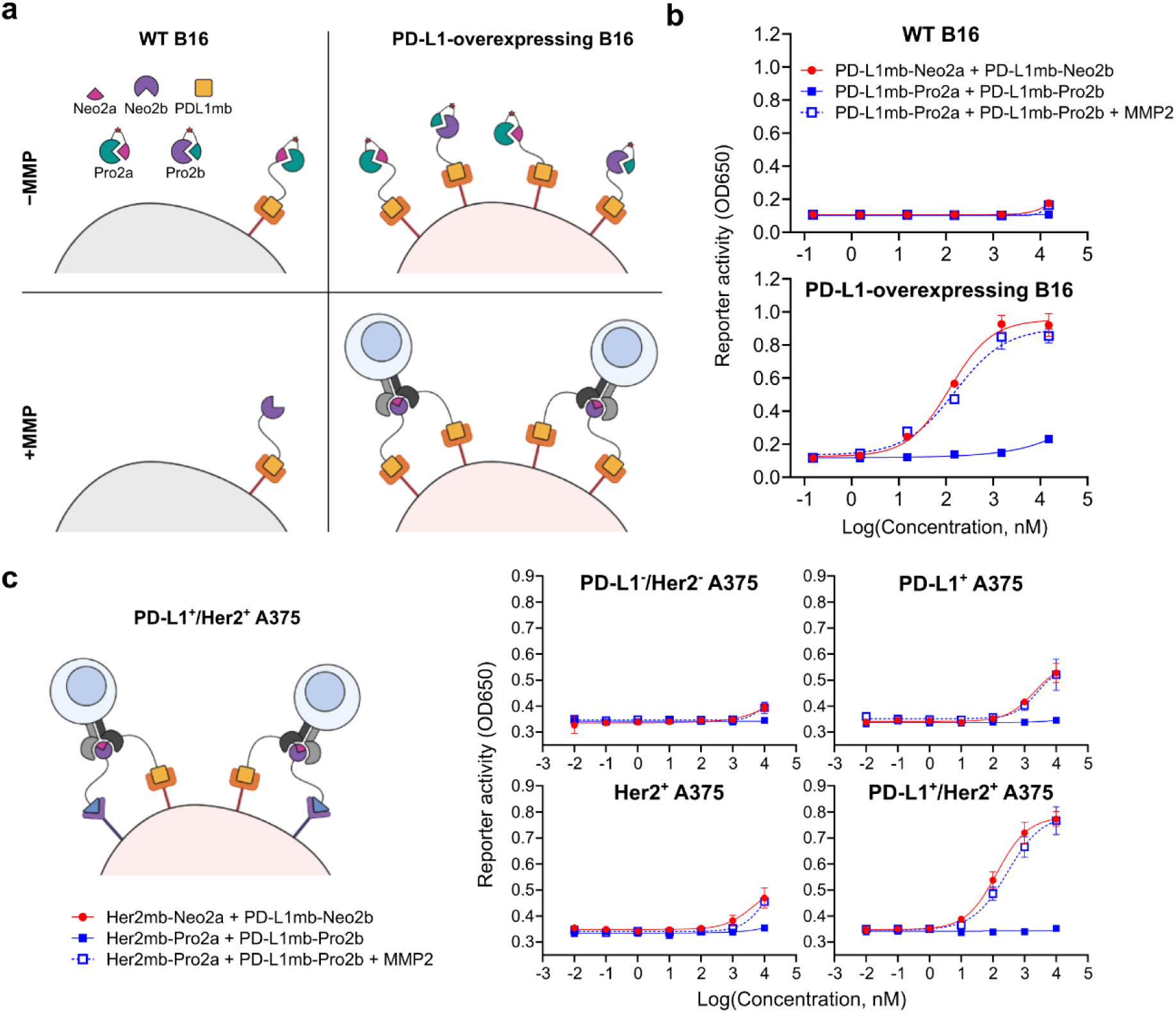
De novo masking enables multi-input control of split Neo2 activity. **a,** Schematic of PD-L1-targeted split Neo2 activation. Targeting domains promote colocalization of split cytokine fragments on PD-L1-expressing cells, while MMP-dependent cleavage removes the masks and permits productive reconstitution of the active cytokine mimic. **b,** Reporter-cell co-culture assays comparing PD-L1-targeted unmasked and masked split Neo2 on wild-type B16 cells or PD-L1-overexpressing B16 cells. Reporter activity remained low in wild-type B16 co-cultures under all conditions. In PD-L1-overexpressing B16 co-cultures, the unmasked targeted pair activated reporter cells, whereas the masked pair showed minimal activity without MMP2 treatment and regained activity after MMP2 cleavage. **c,** Dual-antigen split Neo2 activation using PD-L1- and HER2-targeted fragments. Reporter activity remained low on double-neagtive cells, was intermediate on single-positive cells, and was highest on PD-L1^+^/HER2^+^ A375 cells after MMP2 treatment. Data are mean ± s.d. of three technical replicates.

We then extended the same logic to a dual-antigen configuration by fusing the two split components to binders targeting PD-L1 and HER2, respectively. In this format, reporter activation remained low on wild-type cells, increased modestly on single-positive cells, and was strongest on double-positive PD-L1^+^/HER2^+^ cells after MMP2 treatment (Fig. 2c). Thus, de novo masking and targeting-driven colocalization provide a route to an AND-gated Neo2 system controlled by two molecular inputs: proteolytic unmasking and antigen-dependent reconstitution.

### Linker tuning improves the in vivo therapeutic window of Pro21

We next evaluated whether protease-activatable masking could improve the in vivo tolerability of a potent IL-21 mimic while preserving antitumor activity. We first tested Pro21-L1, which incorporated the same protease-cleavable linker used for the initial in vitro characterization of Pro21. In the MC38 colon carcinoma model, 21h10 produced strong tumor control but was accompanied by progressive body-weight loss, consistent with treatment-associated cytokine activity outside the tumor. As we have previously reported, this 21h10-dependent weight loss results from pancreatitis^7^. At a dose equimolar to 21h10, Pro21-L1 reduced treatment-associated body-weight loss but provided limited tumor control. Increasing Pro21-L1 to a 10× dose improved antitumor activity; however, the dose-matched Pro21-NC also showed partial tumor suppression (Fig. S8). Thus, although the initial masked format improved tolerability, the activity of the non-cleavable control suggested that the antitumor effect at the higher dose was not fully dependent on proteolytic cleavage.

We hypothesized that tuning the sensitivity of the protease-cleavable linker could improve cleavage-dependent control of antitumor activity. A linker that is cleaved too slowly may limit activation in the tumor, whereas a linker that is cleaved too readily could increase activation outside the tumor and narrow the tolerability window. To test this, we generated a series of MMP-cleavable linker variants and compared their sensitivity to MMP2 by enzyme titration (Table S1). Linker exchange did not substantially alter Pro21 masking in cell-based signaling assays: all variants showed similar attenuation in the intact state and comparable recovery after MMP2 treatment (Fig. S9). However, the variants differed in MMP2 sensitivity, with an apparent ranking of L3 > L2 > L4 > L1 (Fig. S10). In MC38 tumor-bearing mice, the more sensitive Pro21-L2 and Pro21-L3 constructs suppressed tumor growth but were associated with treatment-related toxicity evident by progressive body-weight loss, particularly for Pro21-L3 (Fig. S11). These results suggested that maximal linker-cleavage sensitivity is not necessarily optimal in vivo and led us to select the intermediate-sensitivity Pro21-L4 linker for further evaluation.

In MC38 tumor-bearing mice, Pro21-L4 produced dose-dependent tumor control at 2× and 5× molar doses relative to 21h10 (Fig. 3a). At the 5× dose, Pro21-L4 produced sustained tumor control and improved survival, whereas dose-matched Pro21-NC showed reduced antitumor activity (Fig. 3a,b). 21h10 also suppressed tumor growth but was associated with substantial toxicity-related body-weight loss, while Pro21-L4 maintained body weight at both doses (Fig. 3c).

**Figure 3.**
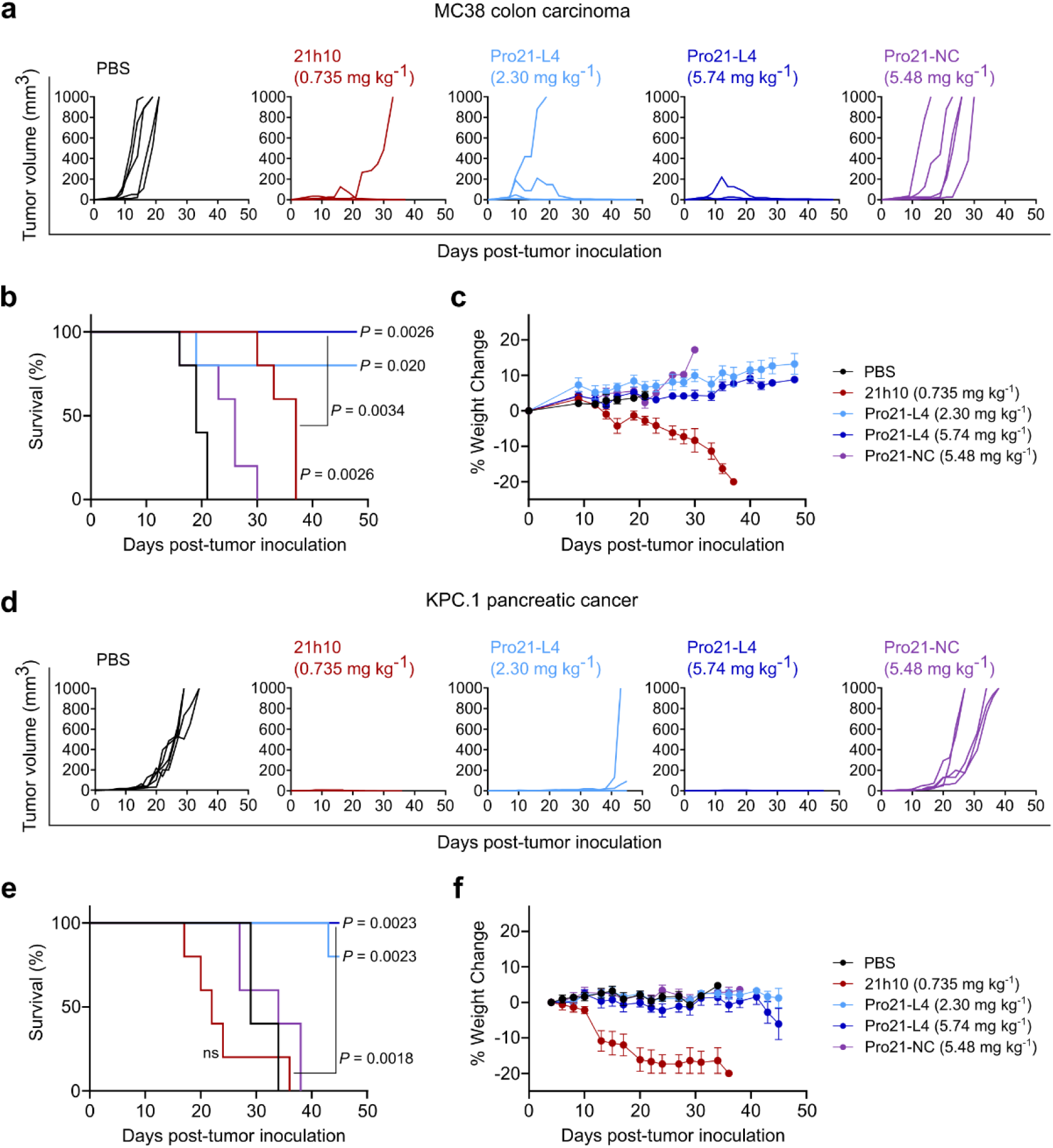
Pro21 retains antitumor activity while reducing treatment-associated body-weight loss in vivo. **a,** Individual tumor growth curves for MC38 colon carcinoma-bearing mice treated with PBS, 21h10, Pro21-L4 at two dose levels, or Pro21-NC. **b,c,** Kaplan–Meier survival analysis (**b**) and percent body-weight change (**c**) in MC38 tumor-bearing mice. **d,** Individual tumor growth curves for KPC.1 pancreatic cancer-bearing mice treated with PBS, 21h10, Pro21-L4 at two dose levels, or Pro21-NC. **e,f,** Kaplan–Meier survival analysis (**e**) and percent body-weight change (**f**) in KPC.1 tumor-bearing mice. n = 5 mice per group. Body-weight data are shown as mean ± s.e.m. *P* values are indicated on the plots; ns, not significant.

We then tested Pro21-L4 in the KPC.1 pancreatic cancer model, where it again suppressed tumor growth and improved survival while avoiding the degree of toxicity observed with 21h10 (Fig. 3d-f). In both models, the lower activity of the non-cleavable control relative to Pro21-L4 supported a contribution of proteolytic unmasking to efficacy. Together, these data indicate that Pro21 retains antitumor activity while reducing treatment-associated weight loss, and that optimizing its linker-cleavage sensitivity can further improve its therapeutic window.

### PD-L1 targeting increases Pro21 activity at reduced dose while preserving evidence of protease-dependent control

Because untargeted Pro21-L4 required a higher molar dose than 21h10 to achieve robust tumor control, we next investigated whether target-mediated localization could preserve antitumor activity at a lower administered dose by concentrating the drug on PD-L1-expressing cells, which are abundant in the tumor microenvironment^22^. We therefore evaluated PD-L1-targeted Pro21-L4 in the KPC.1 model at a dose equimolar to 21h10.

Untargeted 21h10 and PD-L1-targeted 21h10 both produced strong tumor control (Fig. 4a). Although PD-L1 targeting delayed the onset of toxicity-related body-weight loss and extended survival relative to untargeted 21h10, PD-L1-targeted 21h10 was still associated with substantial body-weight loss, indicating that targeting mitigated but did not fully eliminate treatment-associated toxicity from the cytokine (Fig. 4b,c). In contrast, PD-L1-targeted Pro21-L4 produced sustained tumor control and improved survival without appreciable toxicity, whereas the non-cleavable targeted control showed reduced antitumor activity (Fig. 4a-c).

**Figure 4.**
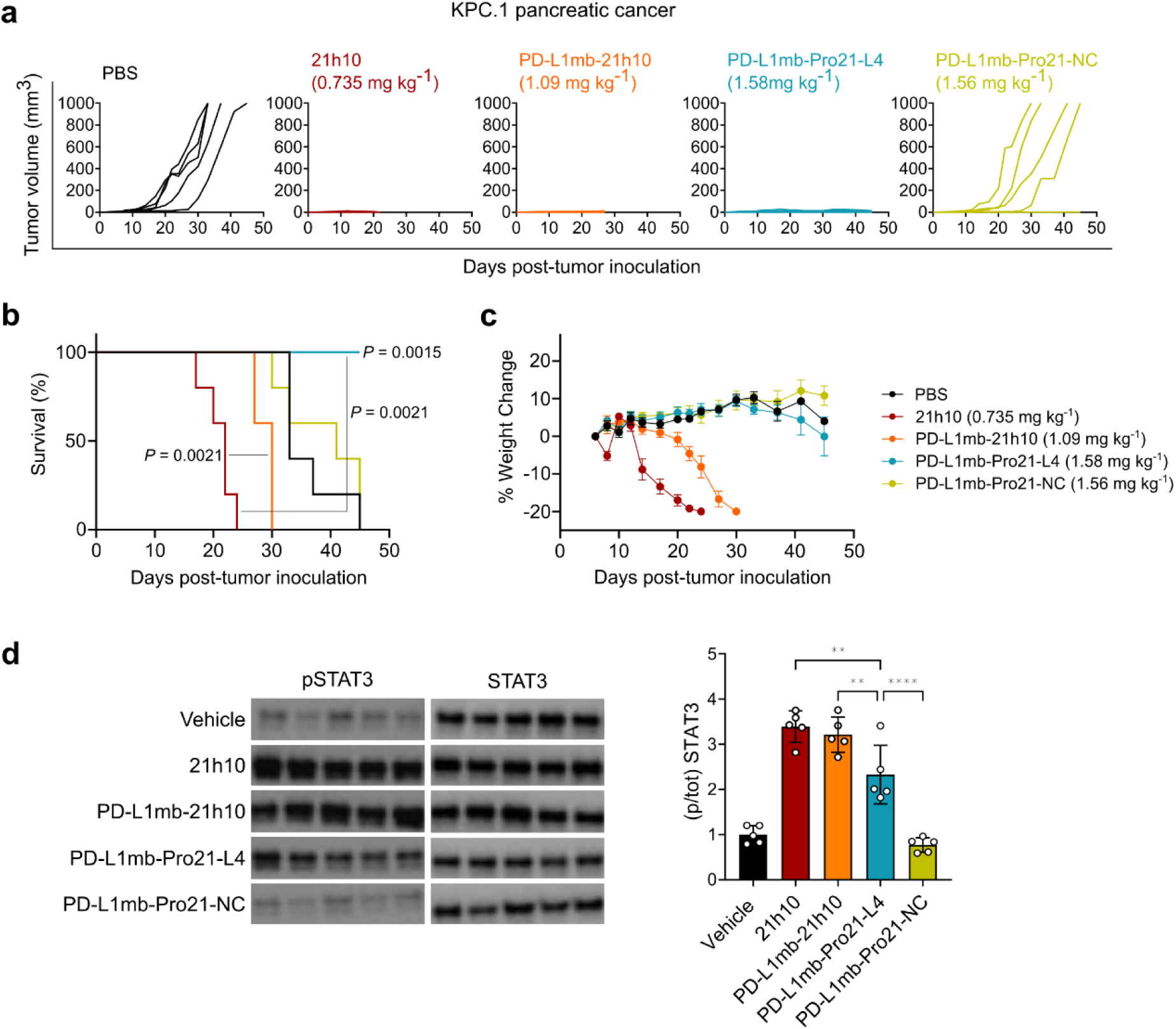
PD-L1 targeting enhances the therapeutic window of protease-activatable Pro21 in KPC.1 tumors. **a,** Individual tumor growth curves for KPC.1 pancreatic cancer-bearing mice treated with PBS, 21h10, PD-L1-targeted 21h10, PD-L1-targeted Pro21-L4, or PD-L1-targeted Pro21-NC. PD-L1-targeted Pro21-L4 maintained strong tumor control at a 1× molar-equivalent dose relative to 21h10, whereas the non-cleavable control showed reduced antitumor activity. **b,c,** Kaplan–Meier survival analysis (**b**) and percent body-weight change (**c**). 21h10 caused rapid body-weight loss, and PD-L1-targeted 21h10 delayed but did not eliminate body-weight loss. In contrast, PD-L1-targeted Pro21-L4 maintained body weight while preserving antitumor activity. n = 5 mice per group. Body-weight data are shown as mean ± s.e.m. *P* values are indicated on the plots. **d,** Mice bearing subcutaneous KPC.1 tumors were treated intravenously with one dose of each of the indicated treatments. Spleens were harvested 3 hours later, lysed in phosphatase inhibitor-containing lysis buffer, and analyzed by immunoblot for pSTAT3 and total STAT3. The ratio of pSTAT3 to STAT3 was quantified. n = 5 mice per group; each lane represents an individual mouse. Full immunoblot images are shown in Fig. S12. \*\**P* < 0.01; \*\*\*\**P* < 0.0001.

To assess peripheral pharmacodynamic activity, we measured STAT3 phosphorylation in the spleen after treatment. PD-L1-targeted Pro21-L4 elicited significantly less STAT3 activation than both untargeted and PD-L1-targeted 21h10 (Fig. 4d). These data indicate that protease-responsive masking suppresses peripheral cytokine signaling under conditions in which tumor targeting alone does not fully restrict activity.

## Discussion

Our results establish de novo masking as a strategy for converting cytokine mimics into protease-activatable prodrugs. De novo masks substantially attenuated receptor binding and signaling while allowing potent activity to be restored by proteolytic cleavage. This approach was effective for both IL-21- and IL-2-like cytokine mimics, supporting its applicability across distinct cytokine receptor systems. A key advantage of de novo masking is that the masking domain is designed directly against the cytokine mimic rather than borrowed from a native receptor, enabling its size, binding geometry, and masking strength to be optimized as intrinsic design parameters. Indeed, we iterated on both the affinity of the masking domain and the protease sensitivity of the linker region several times in the course of this project; rapid design and optimization was critical to success.

Our approach overcomes challenges faced by current tumor-protease-responsive biologics development efforts. Early efforts focused on masked antibody-based formats^23–26^, but this approach has the disadvantage that given the long circulation time of antibodies considerable systemic exposure can occur after mask removal. Masked cytokines^27–29^ and T-cell engagers^30^ are being developed in settings where localized activation could provide substantial benefit but also requires tighter control, as even low levels of systemic activity can drive immune activation or toxicity, requiring a careful balance between systemic suppression and tumor-localized activation. Previously reported protease-activated cytokine prodrugs have generally shown activity shifts in the low- to mid-tens-fold range, with only a subset approaching 100-fold in cell-based assays^11–13,27^. By comparison, Pro21 and Pro2 showed approximately 197- and 150-fold shifts between the intact and protease-cleaved states, respectively, while masking of split Neo2 produced an activity window of at least 800-fold. The compact de novo masks are readily tunable without substantially compromising masking or protease-dependent recovery, enabling iterative optimization toward an in vivo-optimized cytokine prodrug. In the Pro21 series, linker exchange preserved masking and recovery in vitro but altered efficacy and tolerability in vivo, allowing Pro21-L4 to be selected for the best separation between tumor control and body-weight loss. The optimized PD-L1-targeted Pro21-L4 construct retained strong activity even in pancreatic ductal adenocarcinoma, a tumor type that remains largely refractory to most immunotherapies because of dense stroma, poor T-cell infiltration, and an immunosuppressive myeloid-rich microenvironment^31^. Together with its improved tolerability, these results suggest that iterative de novo optimization can produce cytokine prodrugs with the potency and spatial control required for difficult-to-treat tumors.

The masked split Neo2 system extends the platform beyond single-input prodrug activation. By adding antigen-dependent colocalization of complementary cytokine fragments as a second requirement for activation, the masked split architecture converts protease-responsive cytokine prodrugs into an AND-gated system. This layered design could further restrict cytokine signaling to sites where both protease activity and target-antigen expression are present, reducing activity in settings where either cue occurs alone. The ability to combine de novo masks, split cytokine fragments, and tumor-targeting domains suggests a path toward programmable immune agonists that integrate multiple tumor-associated cues.

More broadly, this work illustrates how protein design can be used not only to create potent therapeutic proteins, but also to program when and where their activity is released. Although demonstrated here with de novo-designed cytokine mimics, the same principle should be applicable to a broader range of clinically relevant biologics whose activity is difficult to restrict systemically, including native cytokines and engineered cytokine agonists, antibodies, and T cell-engagers. By converting masking from a fixed inhibitory module into an optimizable design variable, de novo protein design provides a framework for building therapeutic proteins with tunable activity, localization and therapeutic window.

## Methods

### Design of de novo masking domains

Designed masking domains were generated to occlude the receptor-binding surfaces of cytokine mimics. Receptor-bound structures of 21h10 and Neo2 were used to define the target masking interfaces (PDB IDs 9E2T and 6DG5, respectively). RFdiffusion2 was used to generate de novo protein backbones positioned over the receptor-binding surface of each cytokine mimic. The resulting backbones were sequence-designed with ProteinMPNN. Designed mask–cytokine complexes were evaluated using AlphaFold2 (AF2) initial guess predictions. Designs were filtered on predicted structural confidence, mask–cytokine interface confidence and maintenance of the intended blocking geometry. Designs with low mask–cytokine interface predicted aligned error (PAE), high predicted local distance difference test (pLDDT) scores, and no substantial distortion of the cytokine mimic were selected for experimental testing. Candidates were screened by biolayer interferometry (BLI) and HEK-Blue reporter cell assays.

### Interface-focused redesign of masking domains

To improve interface packing, selected mask backbones were subjected to a second round of sequence optimization focused on the mask–cytokine interface. The backbone geometry was held fixed, and residues contacting the cytokine mimic were resampled with ProteinMPNN. Non-interface residues were kept fixed unless local redesign was required to maintain packing. For each selected backbone, redesigned sequences were generated and re-evaluated by AF2. Final designs were selected on the basis of improved interface confidence, preservation of the masked conformation and compatibility with fusion through a protease-cleavable linker.

### Design of masking domains for split cytokine

Masking domains were designed using the same workflow used for intact 21h10 and Neo2 masking domains. Final masking domains were fused to the corresponding split Neo2 components through protease-cleavable linkers. Matched unmasked split fragments and non-cleavable masked controls were generated for comparison. Candidates were screened using HEK-Blue reporter cell assays.

### Protein expression and purification

Genes encoding cytokine mimics and Prokines were codon optimized for Escherichia coli expression and cloned into the LM670 bacterial expression vector with a hexahistidine tag for purification. Proteins were expressed in E. coli BL21(DE3) cells. Cultures were grown in an autoinduction medium supplemented with kanamycin at 37°C until harvest. Cells were collected by centrifugation, and cell pellets were resuspended in lysis buffer (25 mM Tris-HCl, pH 8.0, 300 mM NaCl, 20 mM imidazole, DNase I and protease inhibitors). Cells were lysed by sonication, and lysates were clarified by centrifugation. Clarified lysates were filtered through 0.22-µm filters and applied to Ni-NTA resin equilibrated in the lysis buffer. Bound proteins were washed with wash buffer (25 mM Tris-HCl, pH 8.0, 300 mM NaCl, 20 mM imidazole) and eluted with elution buffer (25 mM Tris-HCl, pH 8.0, 300 mM NaCl, 400 mM imidazole).

Eluted proteins were further purified by size-exclusion chromatography using a Superdex 75 column equilibrated in the SEC buffer (25 mM Tris-HCl, pH 8.0, 300 mM NaCl). Fractions corresponding to the monodisperse peak were pooled and concentrated using centrifugal filters with appropriate molecular-weight cutoffs. Protein purity and apparent molecular weight were assessed by SDS–PAGE, and monodispersity was evaluated from the size-exclusion chromatography profile. Protein concentrations were determined by absorbance at 280 nm using calculated extinction coefficients.

For proteins used in animal studies, low-endotoxin purification procedures were used. Following affinity purification, samples were washed extensively with CHAPS-containing buffer and buffer-exchanged by preparative size-exclusion chromatography using columns precleaned sequentially with NaOH and CHAPS-containing buffer. Final protein preparations were sterile-filtered, concentrated and tested for endotoxin using a limulus amebocyte lysate (LAL) assay. Only preparations below the acceptable endotoxin threshold were used for in vivo experiments.

### Circular Dichroism

Circular dichroism spectroscopy was used to assess secondary structure and thermal stability of purified proteins. Proteins were exchanged into PBS (pH 7.4) and diluted to 2 mg mL^−1^. Far-ultraviolet circular dichroism spectra were collected on a JASCO J-1500 spectropolarimeter using a 1-mm pathlength quartz cuvette. Spectra were recorded from 200 to 260 nm. Buffer-only spectra were collected under the same conditions and subtracted from protein spectra.

Thermal unfolding measurements were performed by monitoring ellipticity at 222 nm while increasing the temperature from 25°C to 95°C at 2°C min^−1^. Where indicated, full-wavelength scans were collected before and after thermal ramping to assess reversibility. Apparent melting temperatures were estimated from the midpoint of the unfolding transition when a cooperative transition was observed.

### Surface plasmon resonance

Surface plasmon resonance measurements were performed on a Biacore 8K instrument (Cytiva) to determine receptor-binding affinities. Receptor ectodomains were immobilized on a CAPture chip according to the manufacturer’s instructions. Purified cytokine mimics or masked cytokine mimics were injected over the sensor surface as serial dilutions in HBS-EP+ running buffer at 30 µL min^−1^. Binding responses were double-reference-subtracted using blank flow cells and buffer injections. Data were analyzed using Cytiva evaluation software. For interactions with well-defined association and dissociation phases, sensorgrams were globally fit to a 1:1 Langmuir binding model.

### Protease Cleavage Assays

Protease-dependent cleavage of masked cytokine constructs was evaluated using recombinant MMP2 or MMP9. Recombinant proteases were activated according to the manufacturer’s instructions and diluted into a cleavage buffer (25 mM Tris-HCl, pH 7.5, 300 mM NaCl, 10 mM CaCl_2_, 0.05% (w/v) Brij-35). Purified protein substrates were mixed with activated protease at the indicated enzyme and substrate concentrations and incubated at 37°C for the indicated times. Cleavage products were resolved by SDS–PAGE and visualized by Coomassie staining.

### HEK-Blue Reporter cell assay

HEK-Blue reporter cells responsive to IL-21 (InvivoGen Cat.code hkb-il21) or IL-2 (InvivoGen Cat.code il2-2) signaling were used to measure cytokine activity. Cells were maintained in DMEM according to the manufacturer’s recommendations and plated in 96-well clear-bottom plates at 5 × 10^4^ cells per well. Purified proteins were prepared as serial dilutions in an assay medium and added to cells. For protease-activated conditions, proteins were pretreated with recombinant MMP2 at 37°C for 2 h before addition to cells. Intact and protease-treated samples were tested in parallel. After 24 h, reporter activity was measured using QUANTI-Blue (InvivoGen Cat.code rep-qbs) according to the manufacturer’s instructions. Absorbance was measured at 650 nm using a plate reader. Dose–response curves were fit using a four-parameter logistic model, and EC50 values were calculated from fitted curves.

### Human PBMC signaling assays

Protein samples were prepared as serial dilutions in the assay medium. PBMCs were stimulated for 15 min at 37°C, then fixed and permeabilized using a fixation buffer (BioLegend, catalog no. 420801) and True-Phos Perm Buffer (BioLegend, catalog no. 425401). Cells were stained with antibodies against lineage markers and phosphorylated STAT proteins. IL-21-like signaling was quantified by pSTAT3 staining, and IL-2-like signaling was quantified by pSTAT5 staining in CD3-positive cells. Samples were acquired on an Attune flow cytometer (Thermo Fisher Scientific), and data were analyzed using FlowJo software. EC50 values were determined by fitting dose–response curves with a four-parameter logistic model.

### Tumor-cell co-culture assays

Tumor-cell co-culture assays were used to evaluate target-dependent cytokine activity. Wild-type or antigen-overexpressing tumor cells were plated in 96-well clear-bottom plates at 2 × 10^4^ cells per well and allowed to adhere overnight. Cells were treated with cytokine constructs for 30 min at 4°C. After washing to remove unbound protein, human PBMCs were added. After 15 min, cells were harvested and stained for flow cytometric analysis. Cytokine signaling was quantified by intracellular pSTAT staining in CD3-positive cells.

### Ethical approvals

All animal protocols were approved by the Dana-Farber Cancer Institute Committee on Animal Care (Protocol #14-019, 14-037, 16-015) and are in compliance with the NIH/NCI ethical guidelines for tumor-bearing animals. C57BL/6 mice were purchased from The Jackson Laboratory (Stock #000664). Survival endpoints for tumor studies included palpable tumor measured or estimated to be greater than 1000 mm^3^, ascites, jaundice, weight loss of 20%, or body condition score less than 2. In all cases where cause of death was unclear, mice were subjected to autopsy to confirm the presence of an end-stage tumor.

### Cancer cell lines and cell culture

MC38 murine colon adenocarcinoma cells (gift from Arlene Sharpe) were cultured in DMEM (Gibco 11-965-092) supplemented with 10% heat-inactivated fetal bovine serum (Gibco Cat# A5670701), 1% 10,000 U/ml penicillin-streptomycin (Gibco 15140-051), and 1% 100 mM sodium pyruvate (Gibco 11360-070). All cell lines were trypsinized with 1x 0.25% Trypsin-EDTA (Gibco 25200-056) and split when they reached 85% confluency in the flask to avoid overconfluency and were maintained at 37°C in a humidified incubator kept at 5% CO_2_. KPC.1 murine pancreatic cancer cells were previously reported^22,32^. These cells were cultured in RPMI 1640 (Gibco 11875093) supplemented with 10% heat-inactivated fetal bovine serum (Gibco Cat# A5670701), 1% 100x GlutaMAX (Gibco 35050-061), 1% 10,000 U/ml penicillin-streptomycin (Gibco 15140-051),1% 100mM sodium pyruvate (Gibco 11360-070), 1% 100x MEM NEAA (Gibco 11140-050), and 1% 1M HEPES (Gibco 15630-080). All cells used for in vivo experiments were negative for known murine pathogens and were implanted at >95% viability.

### Subcutaneous murine tumor models

On Day 0, 6-8-week-old female C57BL/6J mice were subcutaneously implanted using a 150 µL injection of tumor cells suspended in 1x HBSS (Gibco 14025-076), with 1 million cells per mouse for MC38 experiments and 200,000 cells per mouse for KPC.1 experiments. For all experiments, mice were randomized after tumor implantation prior to the start of treatment. Beginning on days 5-7 post-tumor implantation, once tumors were palpable, mice received daily intraperitoneal (IP) injections at the dose indicated per experiment. Mice that did not have palpable tumors at the start of dosing were excluded from the experiment. Mice that had tumors that ulcerated before reaching 500 mm^3^ were also excluded. All constructs were aliquoted and snap frozen in liquid nitrogen to be diluted in 1x endotoxin-free Dulbecco’s PBS, without Ca^2+^ & Mg^2+^ (Millipore-Sigma TMS-012-A) to the appropriate concentrations at the time of dosing. Mice were continuously monitored for survival, weight change, and tumor growth. Endpoint criteria for these experiments were tumor ulceration or tumor volume of 1000 mm^3^ (measured with a ruler, volume calculated using modified ellipsoid formula) or loss of 20% of the initial body weight.

### Western Blot analysis for pSTAT signaling

C57BL/6J mice were administered an intravenous injection with the corresponding constructs at a single dose equivalent to the in vivo IP regimen. After 3 hours, spleens from these mice were harvested and crushed in radioimmunoprecipitation assay (RIPA) lysis buffer supplemented with protease inhibitors (Millipore-Sigma 11836170001) and phosphatase inhibitors (Cell Signaling 5870S). A bicinchoninic acid assay (Thermo Scientific 23225) was performed to determine the protein concentration of these lysates. Equal amounts of spleen lysates across groups were loaded onto 4-20% SDS-polyacrylamide gels (Bio-Rad Cat. #5671095), which were then transferred to a polyvinylidene difluoride (PVDF) membrane (Bio-Rad Cat. #162-0175) using the Bio-Rad Trans-Blot Turbo Transfer System. Membranes were incubated with detection antibodies diluted in a 3% bovine serum albumin (BSA) (Sigma-Aldrich A7906-500G) in Tris-buffered saline with Tween 20 (TBS-T). The following antibodies were used: Phospho-Stat3 (Cell Signaling 9145S) and Stat3 (Cell Signaling 12640S). After imaging the Western blots, the relative density and size of the bands were quantified using ImageJ; ratios of pSTAT3 to STAT3 for all groups were normalized to the PBS control group.

### Statistics

Statistical analyses were performed using GraphPad Prism. Statistical details and sample sizes are provided in the corresponding figure legends. Survival curves were compared using the log-rank (Mantel–Cox) test. Western blot quantification was analyzed using one-way ANOVA followed by Šídák’s multiple comparisons test. Data are shown as the mean, with individual data points shown where indicated.

## Data availability

This paper does not report original code. Other data generated in this study are available upon request.

## Acknowledgments

This work was supported by the National Institutes of Health through the National Cancer Institute (R01CA240339) and the National Institute of Allergy and Infectious Diseases (R0AI160052). This work was also delivered as part of the Matchmakers team supported by the Cancer Grand Challenges partnership, funded by Cancer Research UK (CGCATF-2023/100008), the National Cancer Institute (1OT2CA297288-01), and the Mark Foundation for Cancer Research. SKD was funded by the Ludwig Center at Harvard, Break Through Cancer, NIH R01AI158488, R01CA303051, and U01 CA274276 and is a Member of the Parker Institute for Cancer Immunotherapy. SKD, BDP and AJA were funded by the Hale Family Center for Pancreatic Cancer Research. LQ and SKD were funded by the Claudia Adams Barr Foundation. LQ was funded by NCI K99CA293136. JHC was funded by the Washington Research Foundation and the Kuni Foundation. MD was funded by R01AI169188 and the Peter and Ann Lambertus Family Foundation. We thank Seema Chugh, Ravina Ashtaputre, and Jaclyn Varga-Wiles for maintenance of the KPC mouse colony.

---

**Supplementary Fig. 1.**
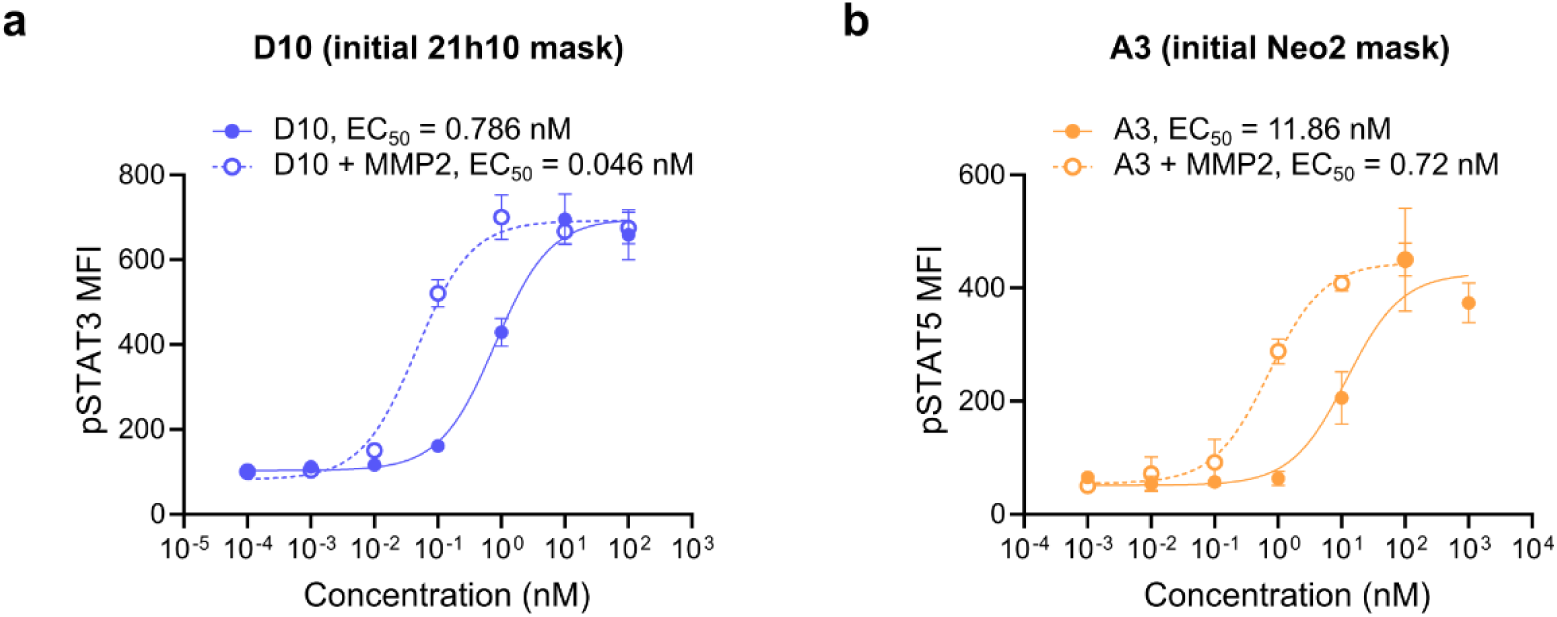
Activity of initial masked IL-21- and IL-2-like cytokine mimics. **a,b,** Human PBMC signaling assays for D10, the initial 21h10 mask (**a**), and A3, the initial Neo2 mask (**b**), before and after MMP2 treatment. D10 and A3 showed attenuated signaling in the intact masked state and increased activity after proteolytic cleavage. pSTAT3 was measured for D10 signaling, and pSTAT5 was measured for A3 signaling. EC50 values are indicated in each panel.

**Supplementary Fig. 2.**
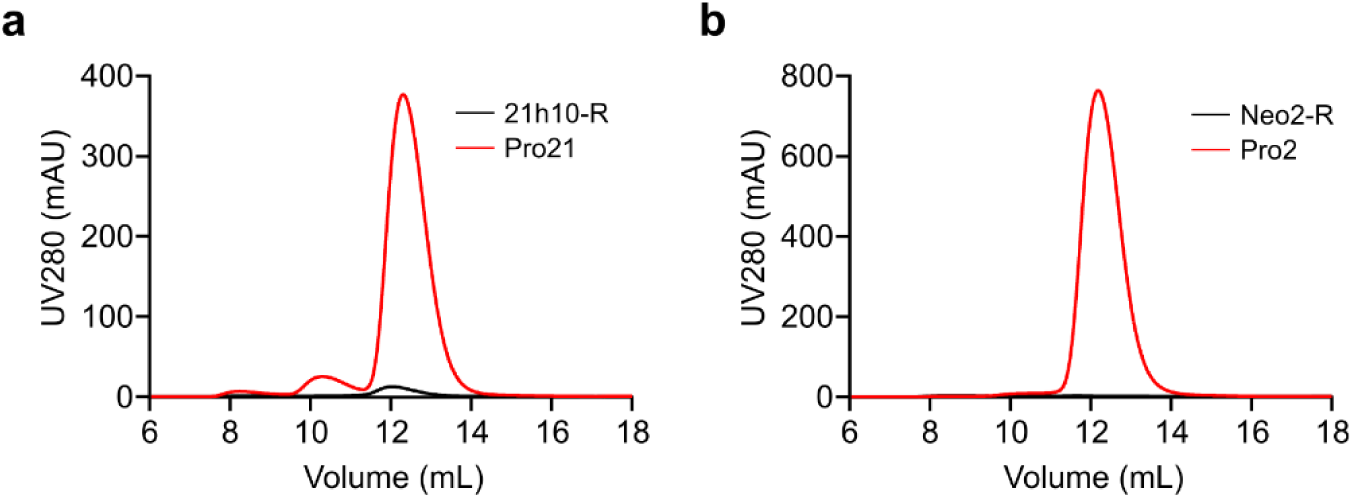
Size-exclusion chromatography analysis of optimized Prokines. **a,b,** Analytical size-exclusion chromatography profiles of purified Pro21 and the corresponding receptor-masked 21h10-R construct (**a**), and Pro2 and the corresponding receptor-masked Neo2-R construct (**b**), monitored by absorbance at 280 nm. Pro21 and Pro2, which contain de novo-designed masking domains, each showed a predominant elution peak, whereas the corresponding receptor-masked constructs showed substantially lower A280 signals under the same production and analysis conditions.

**Supplementary Fig. 3.**
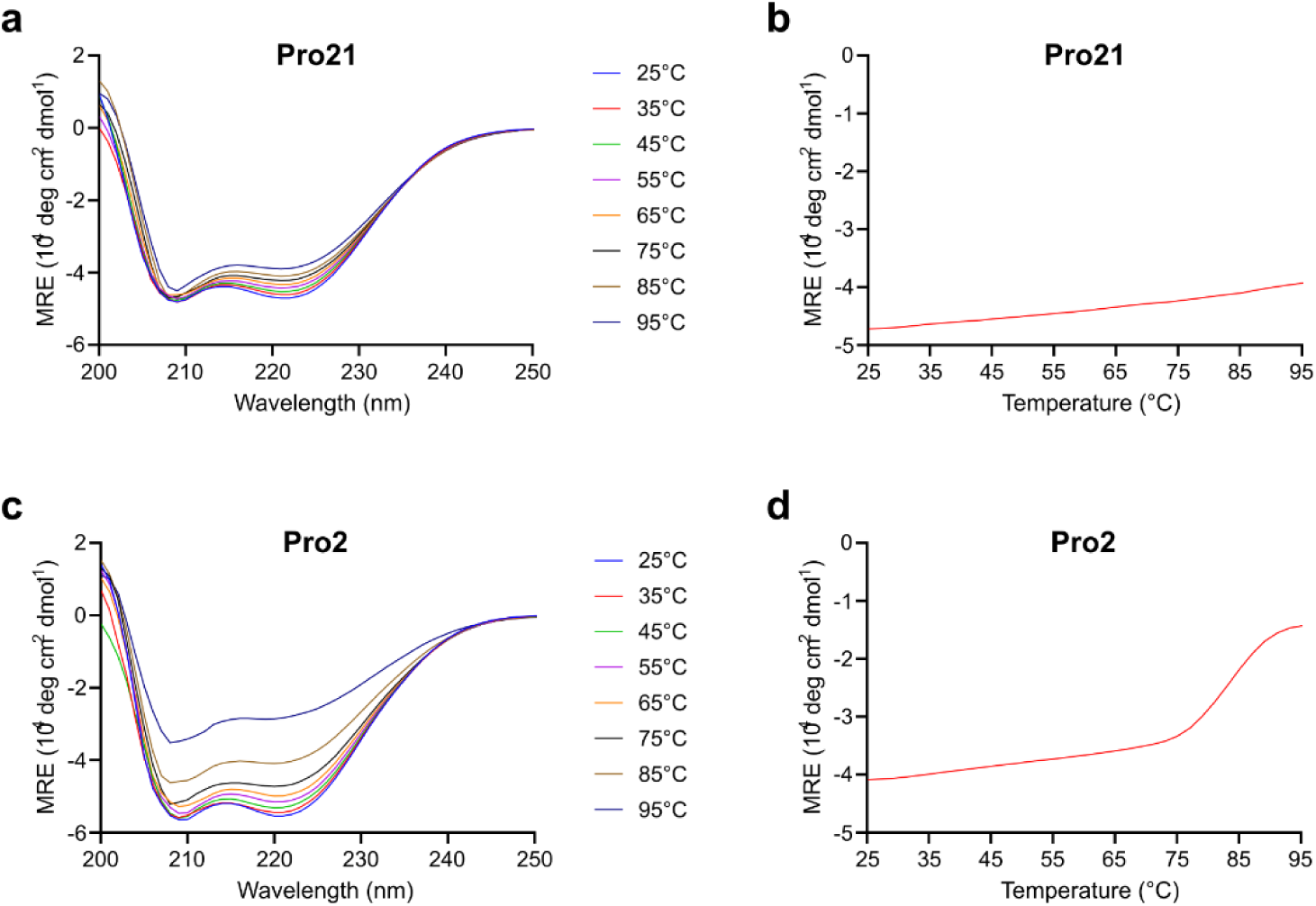
Circular dichroism analysis of Pro21 and Pro2 thermal stability. **a,c,** Far-UV circular dichroism spectra of Pro21 (**a**) and Pro2 (**c**) collected over a temperature range from 25 °C to 95 °C. **b,d,** Temperature-dependent changes in mean residue ellipticity at 222 nm for Pro21 (**b**) and Pro2 (**d**), monitored during thermal ramping.

**Supplementary Fig. 4.**
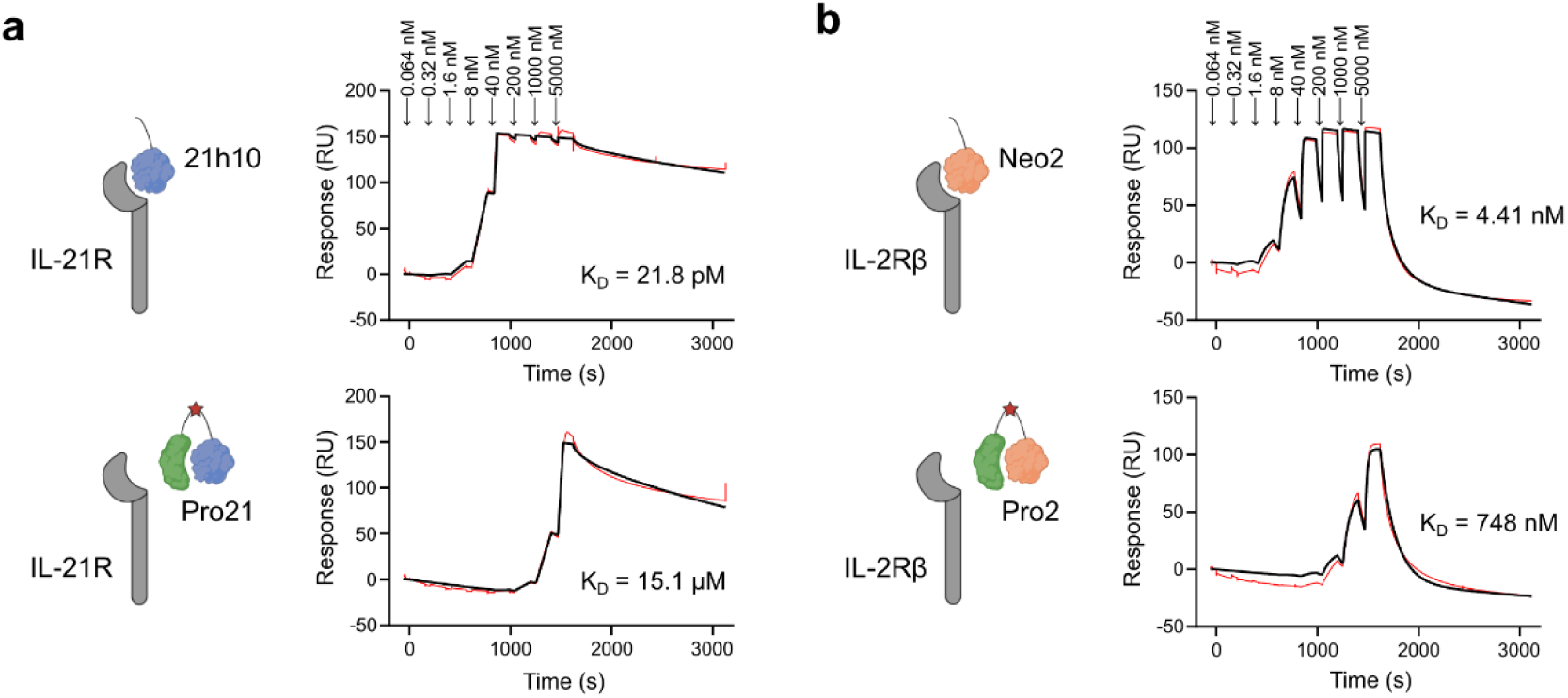
De novo masking reduces receptor binding of Pro21 and Pro2. **a,** Surface plasmon resonance analysis of 21h10 and Pro21 binding to immobilized IL-21R. Masking reduced IL-21R binding affinity from 21.8 pM for 21h10 to 15.1 µM for Pro21. **b,** Surface plasmon resonance analysis of Neo2 and Pro2 binding to immobilized IL-2Rβ. Masking reduced IL-2Rβ binding affinity from 4.41 nM for Neo2 to 748 nM for Pro2. Black traces indicate experimental sensorgrams and red traces indicate fitted binding curves. K_D_ values are indicated in each panel.

**Supplementary Fig. 5.**
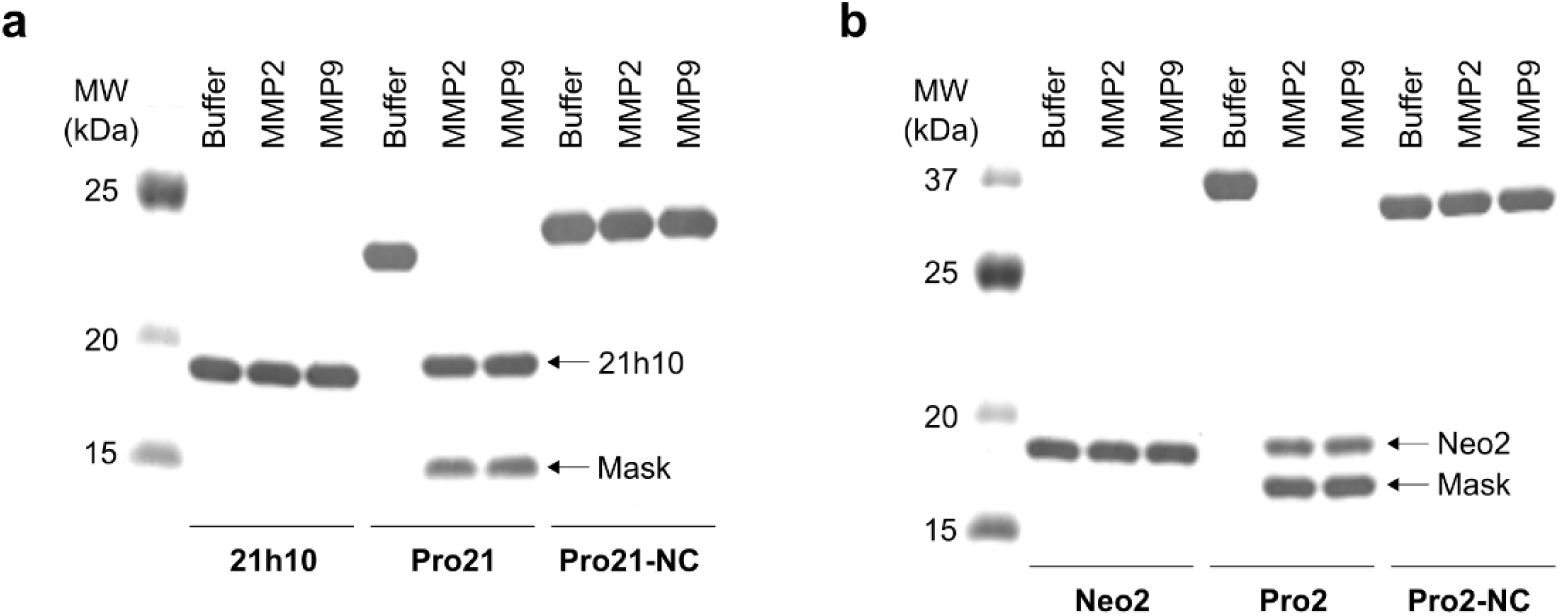
Protease-dependent cleavage of Pro21 and Pro2. **a,** SDS–PAGE analysis of 21h10, Pro21, and Pro21-NC after incubation with buffer, MMP2, or MMP9. MMP2 and MMP9 treatment cleaved Pro21 to generate bands corresponding to 21h10 and the masking domain, whereas Pro21-NC remained intact. **b,** SDS–PAGE analysis of Neo2, Pro2-M, and Pro2-NC after incubation with buffer, MMP2, or MMP9. MMP2 and MMP9 treatments both cleaved Pro2-M to generate bands corresponding to Neo2 and the masking domain, whereas Pro2-NC remained intact.

**Supplementary Fig. 6.**
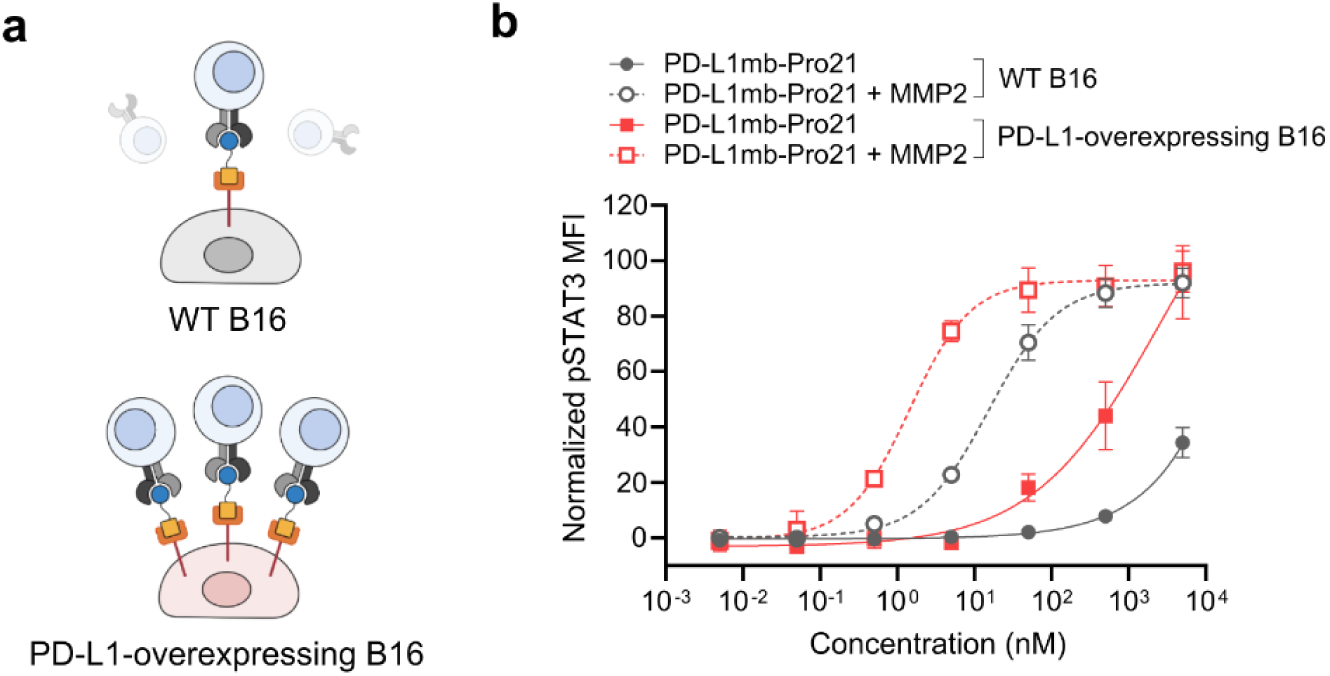
PD-L1 targeting enhances Pro21 activation on PD-L1-expressing cells. **a,** Schematic of co-culture assay comparing PD-L1-targeted Pro21 activity in the presence of wild-type B16 or PD-L1-overexpressing B16 cells. **b,** Human PBMC signaling assay measuring pSTAT3 activation after treatment with PD-L1mb-Pro21 with or without MMP2. PD-L1-overexpressing B16 cells shifted the response of MMP2-treated PD-L1mb-Pro21 to lower concentrations relative to wild-type B16 cells, consistent with PD-L1-dependent enrichment at the target-cell surface. Data are shown as normalized pSTAT3 mean fluorescence intensity (MFI).

**Supplementary Fig. 7.**
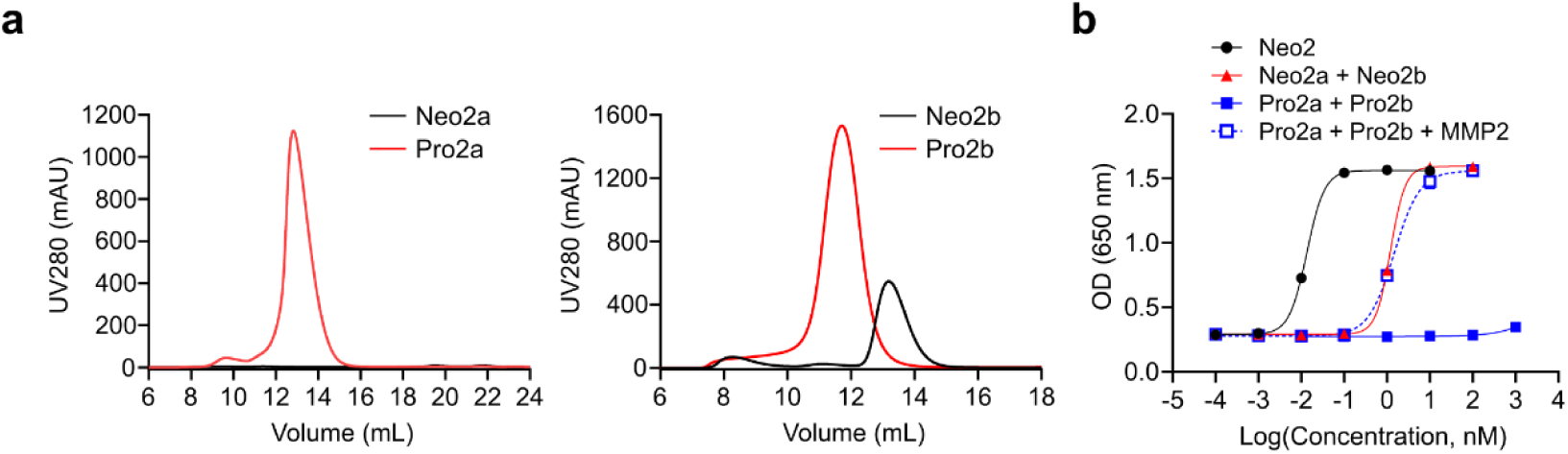
Characterization and protease-dependent activation of masked split Neo2 fragments. **a,** Analytical size-exclusion chromatography profiles of split Neo2 fragments and corresponding masked fragments. Pro2a and Pro2b eluted as predominant soluble species. **b,** HEK-Blue IL-2 reporter assay comparing full-length Neo2, unmasked split Neo2a plus Neo2b, masked Pro2a plus Pro2b, and MMP2-treated Pro2a plus Pro2b. Masked Pro2a and Pro2b showed minimal activity in the intact state, whereas MMP2 treatment restored activity to a level comparable to the unmasked split Neo2 pair.

**Supplementary Fig. 8.**
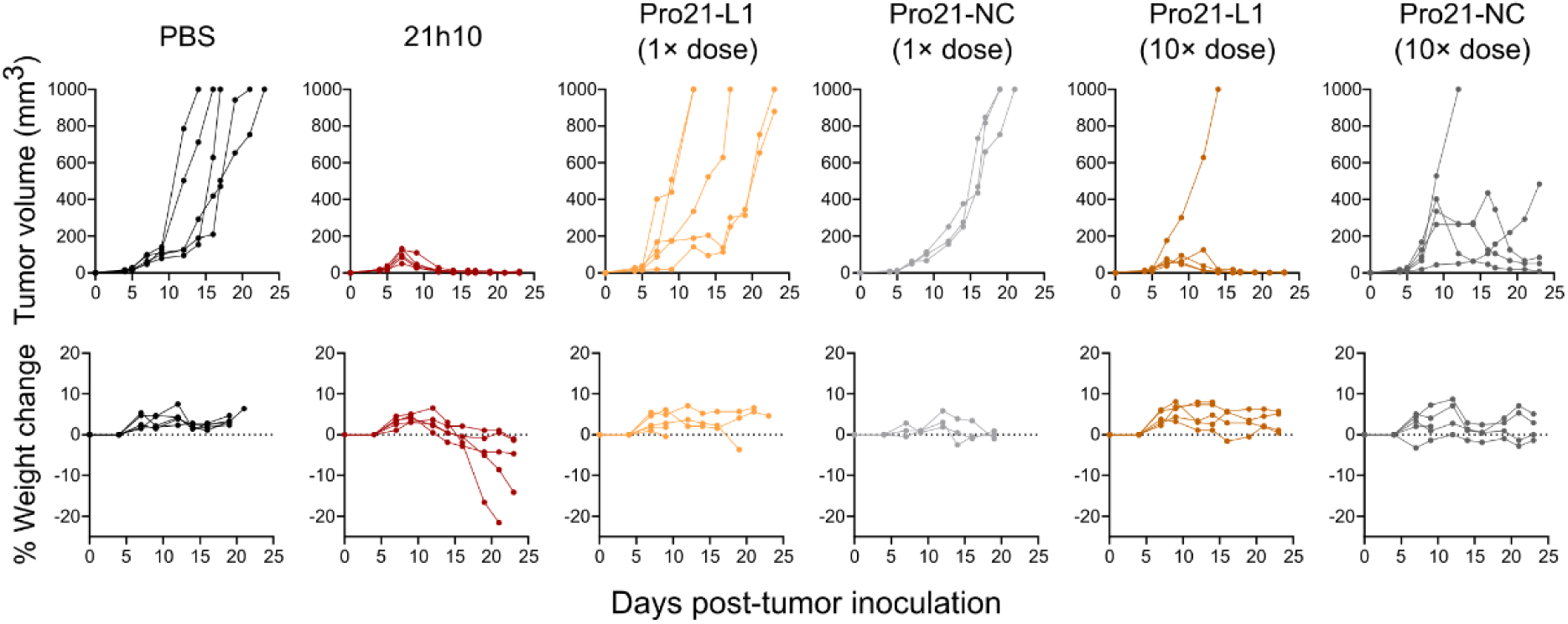
In vivo evaluation of the initial Pro21-L1 construct in the MC38 tumor model. MC38 colon carcinoma-bearing mice were treated with PBS, 21h10, Pro21-L1, or Pro21-NC at 1× or 10× molar-equivalent doses relative to 21h10. Individual tumor growth curves are shown in the top row, and the corresponding percent body-weight changes are shown in the bottom row. 21h10 produced strong tumor control but was associated with body-weight loss. Pro21-L1 reduced body-weight loss relative to 21h10 but showed limited tumor control at the 1× dose, whereas the 10× dose provided greater antitumor activity. The dose-matched Pro21-NC control also showed partial tumor suppression at the 10× dose, suggesting limited cleavage-dependent separation for the initial Pro21-L1 construct under these conditions. n = 5 mice per group.

**Supplementary Fig. 9.**
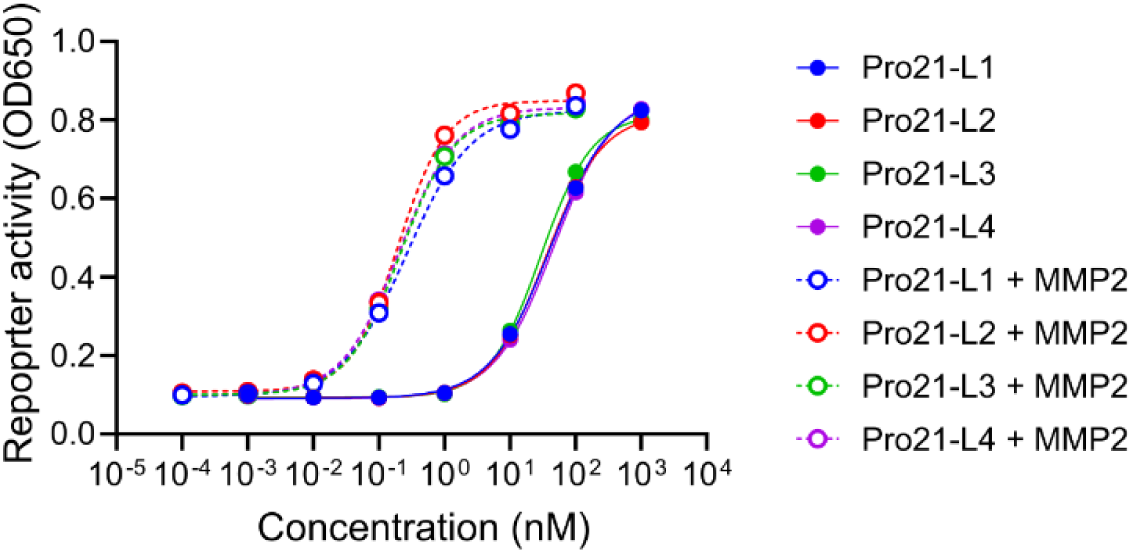
Linker exchange preserves Pro21 masking and protease-dependent recovery. Reporter-cell dose-response assay comparing Pro21-L1, Pro21-L2, Pro21-L3, and Pro21-L4 before and after MMP2 treatment. All linker variants showed similar attenuation in the intact state and comparable recovery of signaling activity following MMP2 treatment, indicating that linker exchange did not substantially alter Pro21 masking or protease-dependent restoration of activity.

**Supplementary Fig. 10.**
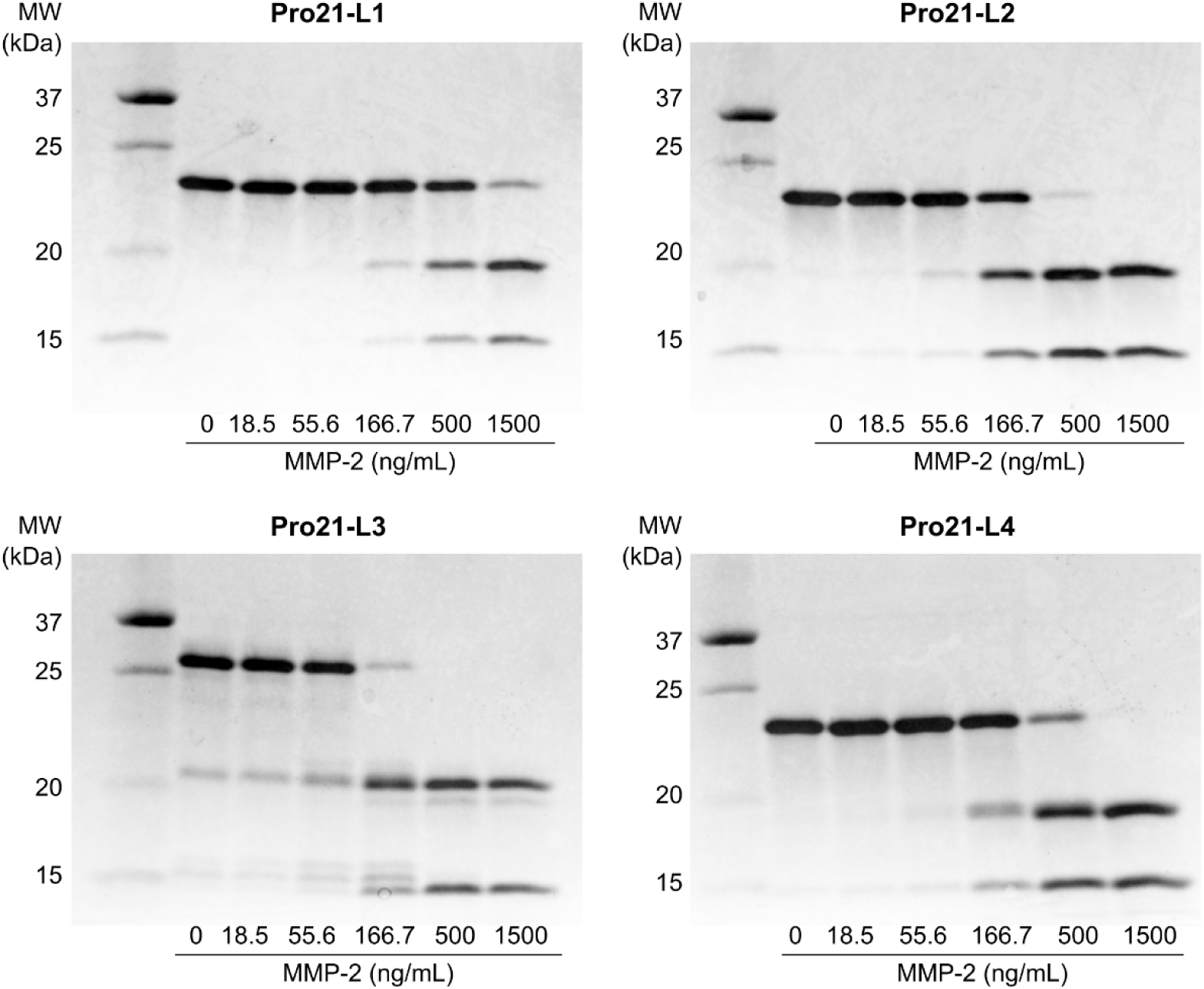
MMP2 sensitivity of Pro21 linker variants. SDS–PAGE analysis of Pro21 linker variants after incubation with increasing concentrations of MMP2. Pro21-L1, Pro21-L2, Pro21-L3, and Pro21-L4 were treated with the indicated concentrations of MMP2, and cleavage was assessed by SDS–PAGE analysis.

**Supplementary Fig. 11.**
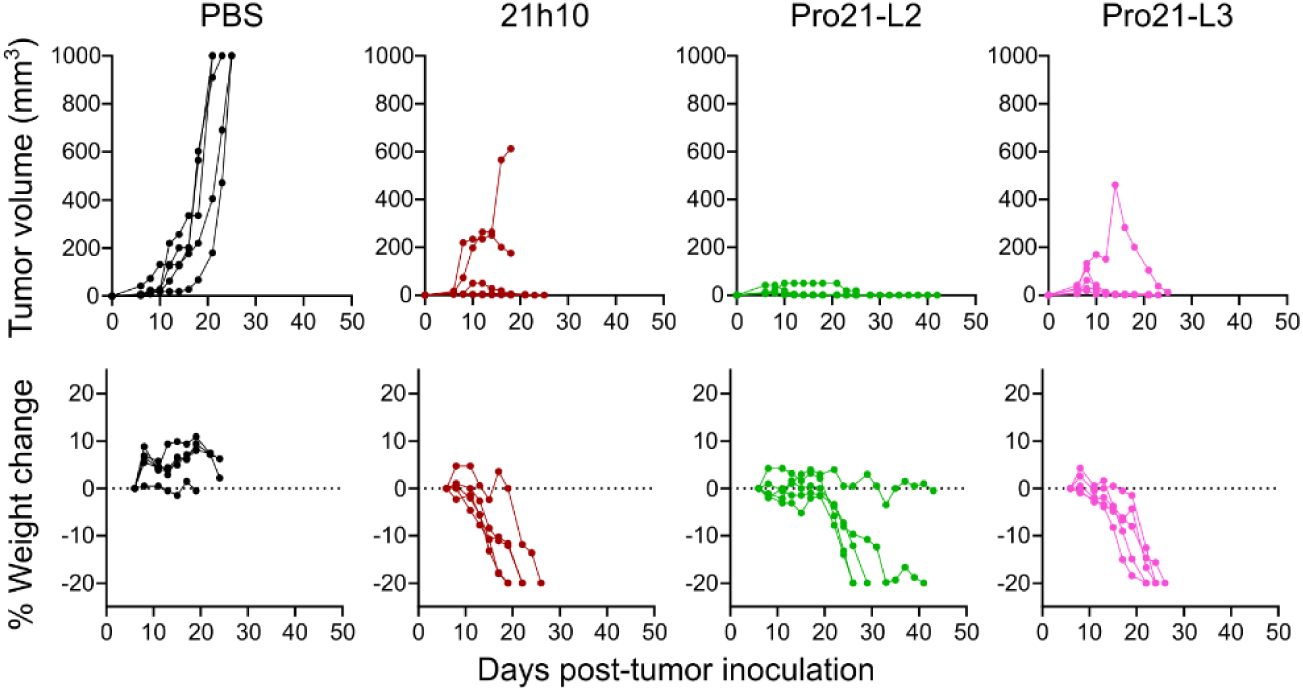
In vivo evaluation of Pro21-L2 and Pro21-L3. MC38 colon carcinoma-bearing mice were treated with PBS, 21h10, Pro21-L2, or Pro21-L3. Individual tumor growth curves are shown in the top row, and the corresponding percent body-weight changes are shown in the bottom row. Pro21-L2 and Pro21-L3 showed strong tumor suppression but were associated with body-weight loss, indicating that increasing linker sensitivity can improve antitumor activity while narrowing the tolerability window. n = 5 mice per group.

**Supplementary Fig. 12.**
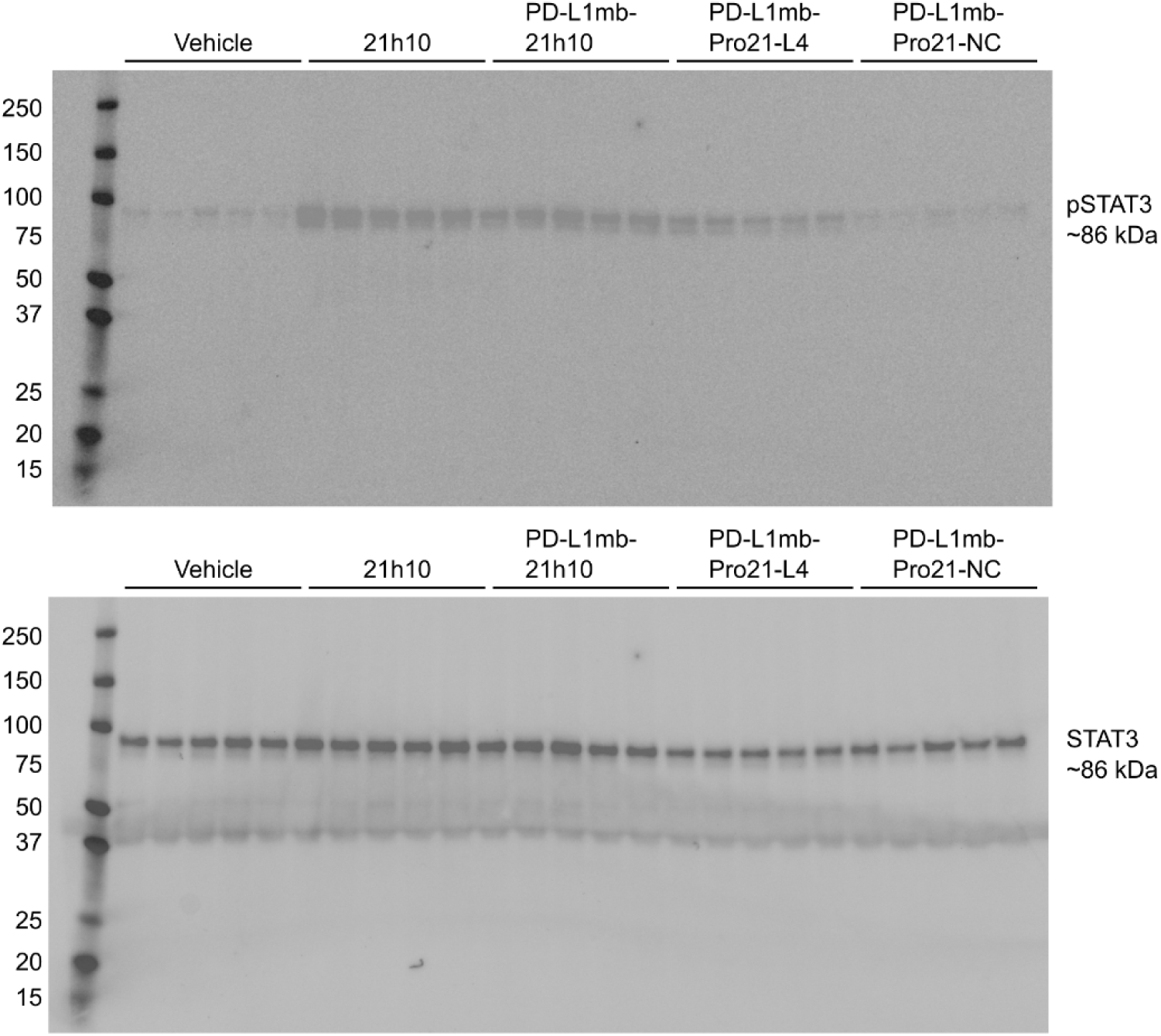
Full immunoblot images for splenic STAT3 signaling. Full immunoblot images corresponding to Fig. 4d showing pSTAT3 and total STAT3 in spleen lysates from the indicated treatment groups. Each lane represents an individual mouse. Molecular-weight markers are indicated in kDa.

**Supplementary Table 1.** Protease-cleavable linker sequences used in Prokine constructs. Sequences are shown from N to C in the orientation used between the cytokine mimic and masking domain. Arrows indicate the expected scissile bonds based on previously reported substrate sequences or motif orientation.

| Linker ID | Linker sequence | Expected cleavage site | Source |
| --- | --- | --- | --- |
| L1 | (GGGS)<sub>2</sub>-(GPLGIAGQ)-(GGGS)<sub>2</sub> | GPLG↓IAGQ | 33 |
| L2 | (GGGS)<sub>2</sub>-(HPVGLLAR)-(GGGS)<sub>2</sub> | HPVG↓LLAR | 12,34 |
| L3 | (GGGS)-(HPVGLLAR)<sub>3</sub>-(GGGS) | HPVG↓LLAR | 12,34 |
| L4 | (GGGS)-(GPLGIAGQ)<sub>2</sub>-(GGGS) | GPLG↓IAGQ | 33 |
| NC | (GGGS)<sub>6</sub> |  |  |

